# Diabetes induces differences in costameric proteins and increases cardiomyocyte stiffness

**DOI:** 10.1101/2024.03.05.583628

**Authors:** G. Romanelli, L. Villarreal, C. Espasandín, J.C. Benech

## Abstract

Several studies demonstrated that Diabetes mellitus can increase the risk of cardiovascular diseases, and remains the most common cause of death in these patients. Cardiomyocytes are subjected to continuous mechanical stress and some proteins like the costamere complex proposed as mechanosensors and mechanotransducers that directly sense and respond to mechanical loads. Costameres sense and transduce it both as lateral force and biochemical signals and are closely related to cardiac physiology because individual heart cells are connected by intercalated discs, which synchronise muscle contraction. Diabetes has an impact on the nano-mechanical properties of living cardiomyocytes, resulting in increased cellular stiffness, as evidenced in clinical studies of these patients and elevated diastolic stiffness. Whether costameric proteins are affected by diabetes in the heart has not currently been studied. In this work, we analyse the effect induced by diabetes in the heart on costameric proteins. The samples (tissue and isolated cardiomyocytes) were analysed by immunotechniques by laser confocal microscopy. Significant statistical differences have been found in the spatial arrangement of the costamere proteins. However, these differences are not due to their expression as evidenced by the Western blot analysis. Heart disease causes an alteration in myocardial relaxation and an increase in left ventricle stiffness, causing a decrease in ejection fraction. Atomic force microscopy was used to compare intrinsic cellular stiffness between diabetic and normal live cardiomyocytes and obtain the first elasticity map sections of diabetic living cardiomyocytes. The data obtained demonstrated that diabetic cardiomyocytes had higher stiffness than control. The present work shows experimental evidence that intracellular changes occur related to cell-cell and cell-extracellular matrix communication which could be related to cardiac pathogenic mechanisms. These changes could contribute to alterations in cardiomyocytes’ mechanical and electrical properties and consequently of the myocardium, producing several cardiac pathologies.

**What Is Known?:**

- Costameres are closely related to cardiac physiology because cardiomyocytes are connected by intercalated discs, which synchronise muscle contraction.
- The structural organisation of the cardiomyocyte proteins is critical for its efficient functioning as a contractile unit in the heart.
- Changes in cellular stiffness are hallmark characteristics of several diseases.

**What New Information Does This Article Contribute?:**

- Whether costameric proteins are affected by diabetes in the heart has not currently been studied. This work shows that T1DM induce significant changes in the spatial organisation of costamere proteins, T-tubules, and intercalated discs.
- The statistical differences in the spatial organisation of costamere proteins were not due to differential protein expression as evidenced by the Western blot analysis.
- Cardiomyocytes are subjected to continuous mechanical stress and some proteins as the costamere complex proposed as mechanosensors and mechanotransducers that directly sense and respond to mechanical loads. Several cardiomyopathies, such as myocardial infarction, are characterised by increased fibrosis and heart stiffness. We obtained the first elasticity map sections (10 μm^2^) of living diabetic cardiomyocytes to compare intrinsic cellular stiffness between diabetic and normal cardiomyocytes. The results show statistical differences in the map sections of T1DM and control cardiomyocytes. T1DM cardiomyocytes are stiffer than normal ones.

## Introduction

Cardiomyocytes can sense and respond to mechanical load through a process named mechanotransduction. Mechanotransduction and its downstream effects function initially as adaptive responses that serve as compensatory mechanisms during adaptation to the initial load^1,2^. Cardiomyocytes are subjected to continuous mechanical stress, and cell-cell junctions support heart contraction and provide electro-mechanical coupling^3^. Several proteins have been proposed as key mechanosensors and mechanotransducers that directly sense and respond to mechanical loads^1^. In 1983, Pardo et al. introduced the name costameres (from Latin costa=rib, and Geek meros=part) to describe a morphological structure in striated muscle^4^. Costameres link myofibrils to the extracellular matrix and sarcomere to the sarcolemma via the Z-disc and M-band^5-7^. These links are closely related to cardiac physiology because individual heart cells are connected by intercalated discs, which synchronise muscle contraction^6^. At the ultrastructural level, it is reported that costameres are closely related to the force necessary for the contraction of the heart muscle by bidirectionally and mechanically linking the cytoskeleton to the extracellular matrix^6,8,9^. Costameres sense a mechanical load and transduce it both as lateral force and biochemical signals^10^. The main components of the costamere are the vinculin-talin-integrin complex (Figure 1), dystroglycan-glycoprotein complex and spectrin complex^11,12^. Kostin et al. found that vinculin, talin and α5β1-integrin are colocalised in the costameres^5^, and it could be expected that defects in costameres would compromise muscle strength directly^9^. Mutations in genes encoding costameric proteins have been identified as causal of cardiac diseases like hypertrophic cardiomyopathies, infiltrative diseases of the heart, and many types of muscular dystrophy^9,13^.

**Figure 1.**
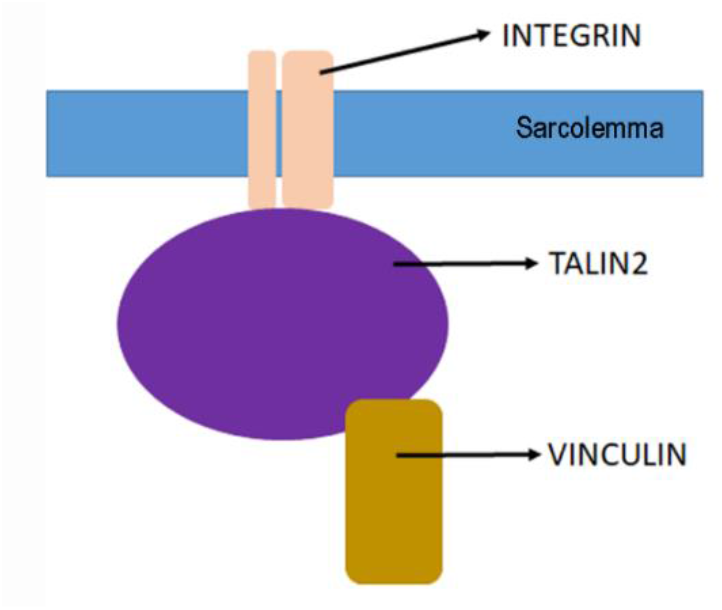
Schematic representation of the vinculin-talin-integrin costameric protein complex in cardiac cells. Graphically assembly of the costamere structure showing the α5β1-integrin, talin 2, and vinculin proteins, based on the work of Zemljic-Harpf et al., 2009. Henderson et al., 2018, 2015, Gorza et al., 2021, and Münch and Abdelilah, 2021.

The sarcolemma and the cell membrane of the cardiac T-tubules were found to contain vinculin, talin, and α5β1-integrin^5^. In the costamere, vinculin is located in the fascia adherens (FA), and FA is part of the intercalated discs. These adhesion sites present roles that link actin filaments to the sarcolemma^6,14^. Talin 2 is expressed at high levels in cardiac muscle and connects integrins to the actin cytoskeleton regulating the affinity of integrins for ligands^15,16^. It has been shown in KO talin 2 mice the importance of this protein in several myopathies^17^. Integrins are a superfamily of glycoproteins that participate mainly in the union of cells with the extracellular matrix, and cell-cell union^18^. Integrins can act as mechanoreceptors^19^, and play important roles in both normal development and the onset of pathologies^18^.

Diabetes mellitus (DM) is one of the most common metabolic disorders in the world and is recognised as a serious public health concern with a considerable impact on human life and health expenditures^20-22^. The incidence of DM in adults has increased over the recent decades. In 2014, 422 million people had DM, and this will increase to 642 million by 2040^21,23^. Diabetes is a major cause of blindness, kidney failure, heart attacks, blood vessels and nerve damage^24^. Several studies demonstrated that DM (hipper and hypoglycemia) can increase the risk of cardiovascular diseases, and remains the most common cause of death in people with DM^25-27^. It is well documented the use of animal models for the study of diabetes. Animal models with Streptozotocin (STZ) administration have been used to induce Type 1 DM (T1DM)^21,28-30^, but there is not much information about the DM effect in the costamere structure.

In 2014, Benech et al. using a T1DM murine model showed a disordered alignment of myocardial cells, cell nuclei of irregular size, fragmented myocardial fibres, and interstitial collagen accumulation The nanoindentation technique with Atomic force microscopy (AFM) was used to evaluate the apparent elastic modulus (AEM) in live isolated cardiomyocytes, both in controls and in diabetic mice. The data obtained demonstrated that diabetes had an impact on the nano-mechanical properties of living cardiomyocytes, resulting in increased cellular stiffness, as evidenced in clinical studies of patients with DM and elevated diastolic stiffness^28^. In 2020, Romanelli et al. used the same animal model to quantificate the spatial organisation of F-actin in striate muscles. The results showed that diabetes has a significant impact on the spatial organisation of F-actin in striated muscles, and these differences were not related to α-actin protein expression according to Western blot analyses^21^. In 2021, Varela et al. obtained the first Young’s modulus images (cellular stiffness) of live cardiac cells (H9c2) by using AFM^31^.

Whether costameric proteins are affected by DM in the heart has not currently been studied. In this work, we analyse the differences in costameric proteins induced by T1DM in the heart in a STZ mouse model. Furthermore, we obtained the first elasticity map sections (10 μm^2^) of living cardiomyocytes to compare intrinsic cellular stiffness between diabetic and normal cardiomyocytes.

## Methods

### Animal care, maintenance and ethical approval

The control mice (n = 12) and T1DM adult male mice (n = 12) were induced as described in Benech et al., 2014 and Romanelli et al, 2020^21,28^. All the mice were weighed, and blood obtained from the tail vein was used for the measurement of nonfasting blood glucose levels using an ACCU-CHEK Compact Plus System (Indianapolis, IN). Every seven days, the blood glucose levels were measured again and the continued diabetic status was monitored. Mice with >250 mg/dL glucose were considered diabetic. The experimental procedures were approved by the Institutional Ethics Committee (CEUA-IIBCE) following the national legislation.

### Samples Processes

Samples for subsequent analyses were obtained from the left heart ventricle of control and diabetic mice. A) The cardiac muscle tissue was cut (10 μm) using a cryostat (MEV, Slee). B) Isolated cardiomyocytes were obtained according to Acker-Johnson et al.^32^. To select viable cardiomyocytes, they were stained with propidium iodide (PI) in a Petri dish chamber. Vital cells in the chamber were selected in an Olympus IX81 inverted microscope and used for atomic force microscopy (AFM). The cardiac tissue and isolated cardiomyocytes were fixated and permeabilised (4% paraformaldehyde + 0.2% Triton X) for laser confocal microscopy (LCM) analysis.

### Histochemistry, cytochemistry and laser confocal microscopy

The samples (tissue and isolated cardiomyocytes) for LCM analysis were incubated with primary antibodies: rabbit monoclonal anti-vinculin (1:75 dilution, ab129002), rabbit monoclonal anti-α5-integrin (1:500, ab179475), mouse monoclonal anti-talin 2 (1:2000, ab105458). The secondary antibodies used were donkey anti-rabbit IgG (1:200, ab96922) for vinculin, goat anti-rabbit IgG (1:250, ab 150077) for integrin αV, and talin-2 antibodies were conjugates with Mix-n-Stain CF555 Dye (92137) following the manufacturer’s instructions. In every case, all the microscope and software settings (laser intensity, high voltage photomultiplier, image depth) were the same for imaging comparing samples. The quantifications on the images of the three proteins were carried out according to the work of Romanelli et al.^21^. To discard that the results were biased by the plane where the images were taken, 10 Z-stacks projections were performed to select the middle plane.

### Analysis of elasticity maps of live cardiomyocytes

Topography and elasticity maps of control mice (n = 9, 20 cells, 10963 force curves) and T1DM mice (n = 8, 27 cells, 12331 force curves) were obtained with an AFM (BioScope Catalyst, Bruker), using silicon nitride probes (DNP-10, Bruker; cantilever D) with a pyramidal tip (nominal radius = 20 nm), attached to a triangular 200-μm-long cantilever (nominal spring constant = 0.06 N/m). After isolation, cardiomyocytes were placed in Tyrode buffer solution (in mM: 135 NaCl, 5.4 KCl, 1 CaCl2, 1 MgCl2, 10 glucose, and 5 HEPES, pH 7.4), and then were plated on glass microslides that were previously treated with polylysine (1x) and laminin (5 μg/ml) to promote cell adhesion to the substrate. The plated cardiomyocytes were placed into an Olympus IX81 inverted microscope coupled to the AFM, whose stage was maintained at 37 ºC. All AFM measurements were conducted within 1 h after plating the cells. Images were obtained using the Peakforce Quantitative Nanomechanical Mapping (PF-QNM) AFM mode, which provides both topographic and mechanical properties maps of the same area at a single scan of the cell surface. The cantilever spring constant was calibrated using the thermal noise method (“thermal tune” application of Bruker AFM software) and was 0.1361 N/m. PF-QNM parameter settings included peak force frequency 0.25 kHz, setpoint 20 nN, amplitude 2000 nm and scan rate 0.2 kHz. For elasticity maps, Derjaguin, Muller, Toropov (DMT) model was used by PF-QNM mode to retrieve Young’s modulus values from the force-indentation curves performed while scanning the sample^33^, given by the equation:

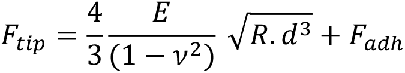

Where Ftip is the force on the tip, Fadh is the adhesion force, R is the tip end radius and d is the tip-sample separation. The Young’s modulus values for the statistical analysis were exported from the elasticity maps of 20 cells from control, and 19 cells from T1DM mice.

### Sample preparation and immunoblotting

Western blot analyses were performed with protein extracts from normal and T1DM mouse tissues. The extracts were prepared from tissue homogenised and lysed with buffer solution: 4-2-Hydroxyethyl-1-piperazine ethane sulfonic acid (HEPES, 20 mM), ethylene glycol tetraacetic acid (EGTA, 1 mM), phenylmethylsulfonyl fluoride (PMSF, 1 mM), 1% Triton X-100, 10% glycerol, Dithiothreitol (DTT, 1mM), sodium fluoride (10 mM), sodium orthovanadate (1 mM), and protease inhibitor cocktail (SigmaFast™Protease Inhibitor Tablets); final concentration 1X; pH 7.4. The protein homogenates were sonicated, and the protein concentration was determined using the Bradford assay. The samples were analysed with denaturing gel electrophoresis (8% and 10% acrylamide). Equivalent protein amounts were loaded into each lane, transferred to a nitrocellulose membrane, and blocked with 5% skim milk powder in Tris-buffered saline containing 1% Triton X-100. The primary antibodies used were rabbit monoclonal anti-vinculin (1:2500 dilution, ab129002), rabbit monoclonal anti-α5-integrin (1:2500, ab179475), mouse monoclonal anti-talin 2 (1:2000, ab105458), and anti-GAPDH (1:5000 dilution, ab181602). The secondary antibody used was goat polyclonal to rabbit IgG Alexa Fluor_®_ 488 (1:1,500 dilution, ab150077) and goat antimouse IgG Alexa Fluor_®_ 488 (1:1,500, ab150113). The signal was developed using a high-performance luminescent image analyzer FLA-9000, according to the manufacturer’s instructions. The bands were quantified using the ImageJ software to determine the relative protein expression levels.

### Quantification and statistical analysis

A total of 150 squares (100 μm^2^) were measured in isolated cardiomyocytes using images from different fields of each mouse sample group. The placement of the squares was such that the tissue occupied the entirety of each square. Images were acquired in the mid-focal plane of the cell sections. Between 100 and 130 images from different fields of each mouse sample were used for the counting and analyses. The analyses of the given samples were independently performed by two researchers, and all the assays were done per triplicate. Regions in which protein signals were over 1.5 μm^2^ were defined as clumps. The protein-occupied area (protein labelled with antibody) of each square was measured as a percentage of the total area. The black spaces occupied by DAPI-stained nuclei were discarded. Intercalated discs per fibre were quantified to obtain the number of intercalated discs of each condition. All the quantifications were done using ImageJ software. Graphs and descriptive and comparative statistical analyses were made using SPSS 10.0. A Shapiro-Wilk test was performed in all comparisons to verify the normality of the data. Mann-Whitney-U, Kolmogorov-Smirnov, Kruskal-Wallis, and Students-t tests (data expressed as medians ± SD) were used for comparing the data of the control and diabetic groups. Values with p < 0.05 were considered statistically significant.

## Results

### Integrin-talin-vinculin differences between control and diabetic mice

Comparisons between costameric complex proteins (Integrin-Talin-Vinculin) in the heart of control and induced diabetic mice after 3 months of STZ injection are shown in Figures 2 to 6. Significant differences were found between control and diabetic mice in the three costameric proteins analysed.

**Figure 2.**
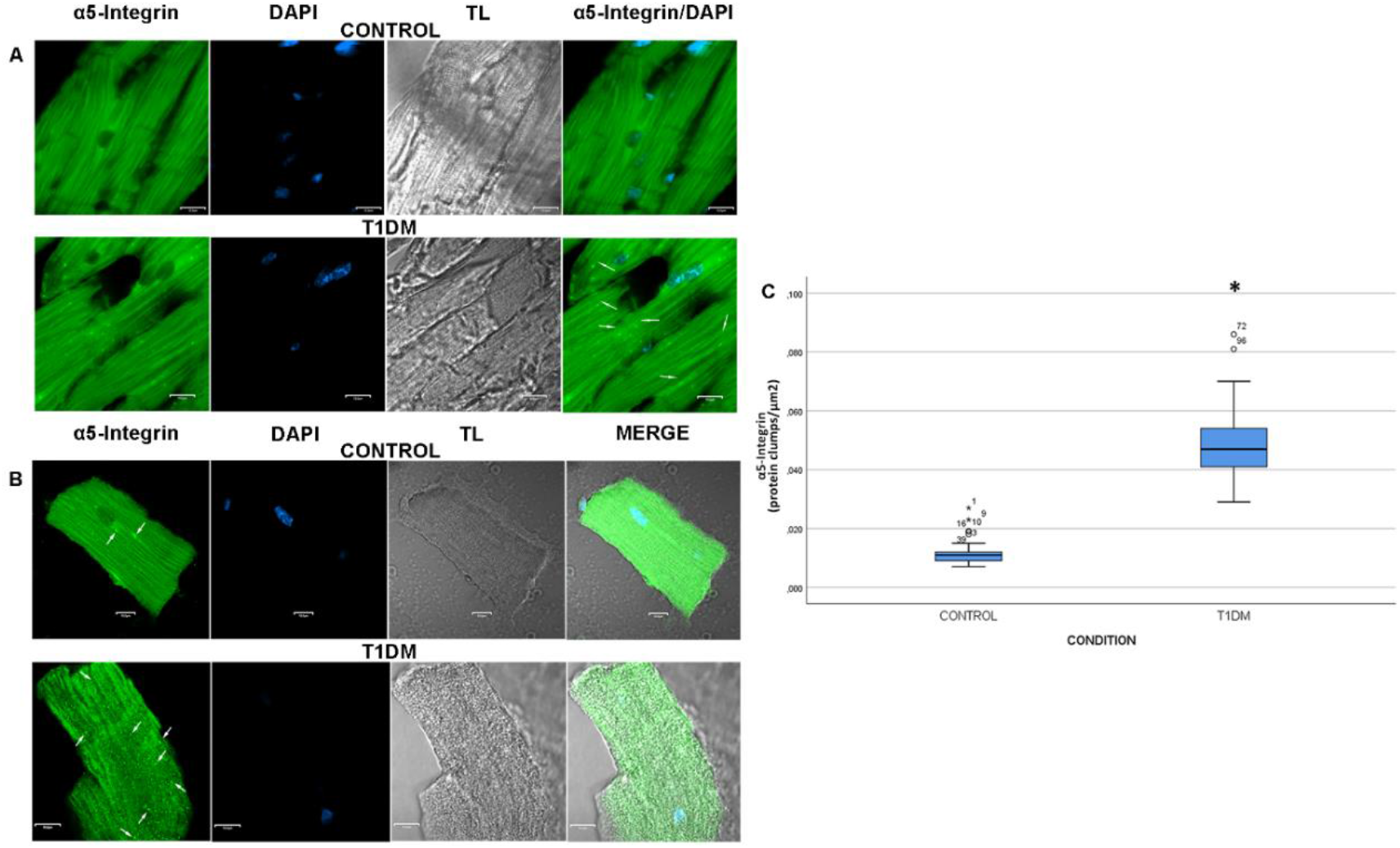
Laser confocal images of representative longitudinal myocardium cross sections and cardiomyocytes of control and diabetic mice labelled for α5-integrin; and statistical analysis. **(A)** Cross sections of the left ventricle of the myocardium show more α5-integrin protein clumps in T1DM (down) than in control mice (up). White arrows sign protein clumps. **(B)** Mice cardiomyocytes show the same pattern as myocardium. Diabetic cardiomyocytes have more α5-integrin protein clumps (arrows) than control ones. **(C)** Mann-Whitney U and Kolmogorov-Smirnov tests (data expressed as medians ± SD) were used for comparing the data of control (n = 8) and T1DM (n = 8) cardiomyocytes (*p < 0.05). There were more protein clumps in diabetic mice than in control ones.

### α5-integrin

As shown in Figure 2A, the myocardium tissue of T1DM mice had more α5-integrin clumps (arrows) than control mice. The same pattern was appreciated in isolated cardiomyocytes (Figure 2B, arrows), and disorganisation in the spatial disposition of this protein was observed. The statistical analysis comparing the number of α5-integrin clumps per area shows significant differences. The diabetic cardiomyocytes had more protein clumps than the control ones (Figure 2C).

### Talin-2

Regarding talin-2 protein, tissue samples also showed more clumps (arrows) in T1DM than in control mice as seen in Figure 3A. At the cardiomyocyte level, the same pattern of clumps can be observed in Figure 3B (arrowheads). The statistical analysis comparing the number of talin-2 clumps per area did not have differences (Figure 3C). However, it can be seen that there is a higher percentage of talin-2 protein unoccupied áreas (Figure 3D) in T1DM cardiomyocytes than in control ones.

**Figure 3.**
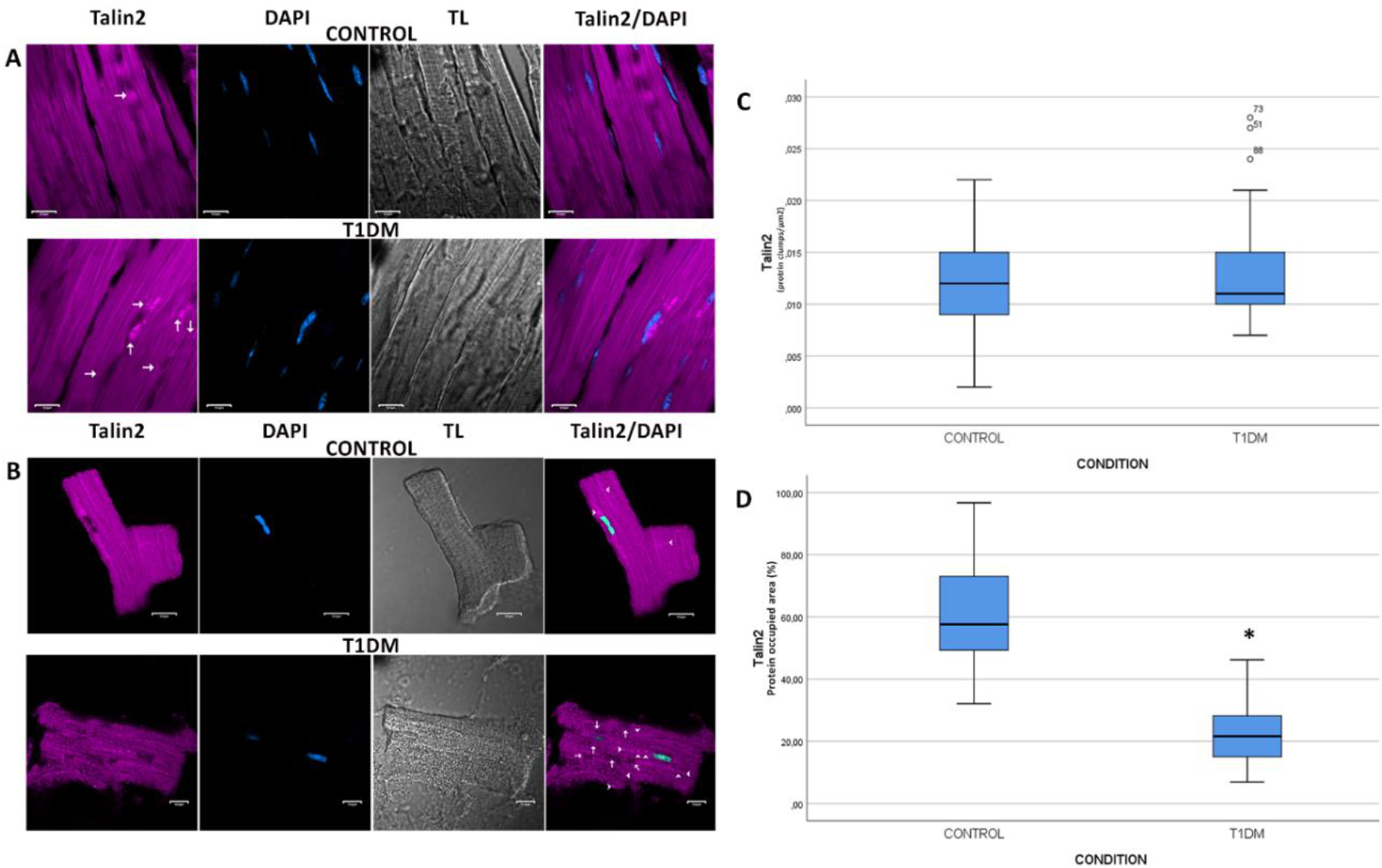
Immunolocalisation of talin 2 protein in longitudinal cross sections of the myocardium and cardiomyocytes of control and T1DM mice. Statistical analysis. **(A)** The representative images of the left ventricle of the myocardium show little more talin 2 protein clumps in T1DM (down) than in control mice (up). Protein clumps are signed by white arrows. **(B)** Cardiomyocyte analysis shows a similar pattern to the myocardium. T1DM cardiomyocytes (down) have more talin 2 protein clumps (head arrows) than control (up) and have more protein unoccupied areas than control ones (arrows). Mann-Whitney U and Kolmogorov-Smirnov tests (data expressed as medians ± SD) were used for comparing the data of control (n = 8) and T1DM (n = 8) mice (*p < 0.05). **(C)** There were no statistical differences between cardiomyocytes of control and diabetic talin 2 protein clumps. **(D)** The talin 2 unoccupied areas were the regions in which talin 2 signals were absent for over 1.5 μm^2^. The DAPI-stained nuclei were discarded. There were statistical differences between control and diabetic cardiomyocytes. Diabetic cardiomyocytes have more talin 2 unoccupied areas than controls. Data are shown as a percentage area of protein signal per total area.

### Vinculin

In cardiac tissue (Figure 4A), control samples showed vinculin signals predominantly in the intercalated discs (arrows) while in T1DM the signals were concentrated on the lateral side of the cardiac fibres (arrowheads). Figure 4B shows that T1DM-isolated cardiomyocytes have more clumps (arrowheads) and more protein-occupied areas (arrows) than control ones. The statistical analysis showed that T1DM had more clumps (Figure 4C), more unoccupied areas (Figure 4D), and less number of intercalated discs (Figure 4E) than controls.

**Figure 4.**
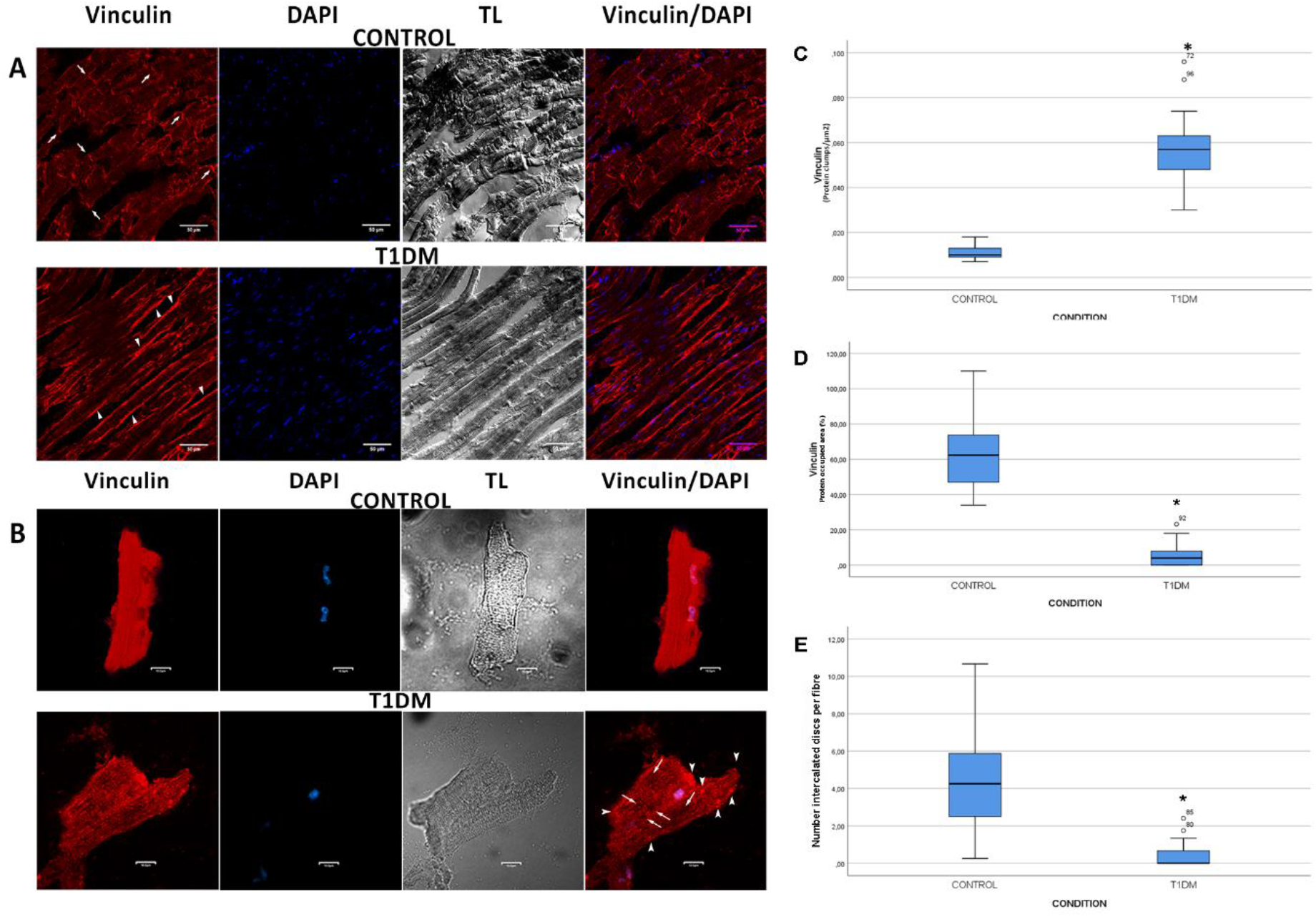
Localisation of vinculin protein in longitudinal cross sections of the myocardium and cardiomyocytes of control and T1DM mice; and statistical analysis of the differences in vinculin protein. **(A)** The left ventricle of the myocardium shows more intercalated discs per muscular fibre labelled with vinculin (arrows) than in diabetic conditions. Additionally, in T1DM left ventricle vinculin protein was laterally located in cardiac fibres (head arrows) **(B)** Representative T1DM cardiomyocytes show more clumps (head arrows) and more vinculin unoccupied areas (arrows) than controls. **(C)** There were statistical differences between control and diabetic vinculin protein clumps. T1DM cardiomyocytes have more vinculin clumps than controls **(D)** The vinculin unoccupied areas were the regions in which vinculin signals were absent for over 1.5 μm^2^. The DAPI-stained nuclei were discarded. There were statistical differences between control and diabetic cardiomyocytes. Control cardiomyocytes have more vinculin-occupied areas than diabetic ones. Data are shown as a percentage area of protein signal per total area. **(E)** The number of intercalated discs per fibre was statistically different between controls and diabetic conditions. Controls have more intercalated discs per fibre than the T1DM condition. Mann-Whitney U and Kolmogorov-Smirnov tests (data expressed as medians ± SD) were used for comparing the data of control (n = 8) and T1DM (n = 8) mice (*p < 0,05).

### Differences in the costamere proteins expression: Western blot analyses

α5-integrin, talin-2 and vinculin expressions in each condition were examined to confirm if the differences observed in the cardiac tissue and cardiomyocytes were a consequence of the protein expression. Specific antibodies against α5-integrin, vinculin and talin-2 were used to analyse both conditions in denaturing gels (8% and 10% respectively). Results show clear bands at 116 kDa (α5-integrin),125 kDa (vinculin), and 272 kDa (talin-2) of the denaturing gel confirming the presence of the three proteins (Figure 5). Glyceraldehyde 3-phosphate dehydrogenase (GAPDH) protein was used as the loading control for the three proteins. A statistical comparison of the three protein expressions between the control and diabetic mice did not show any difference.

**Figure 5.**
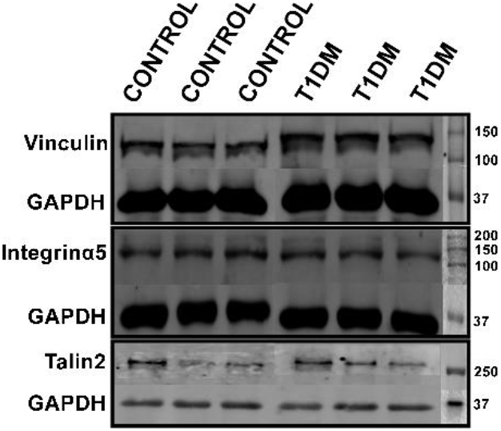
Representative images of Western blots of costamere proteins in control and T1DM STZ injected mice. Protein expressions were quantified relative to GAPDH. There were no statistical differences between the control and T1DM samples.

### Elasticity maps

PF-QNM AFM images included topographic (height and peak force error) and elasticity maps, as displayed in Figure 6A. Topographic maps exhibited myofibrils and cross striations both in control and T1DM mice cardiomyocytes, although T1DM mice cardiomyocyte striations were qualitatively observed as smoother. Elasticity maps revealed a general stiffness increase for T1DM mice cells, confirmed by the statistical analysis of exported Young’s modulus values (presented as the histogram in Figure 6B). The resulting Young’s modulus values, expressed as medians ± SD, were significantly different, as follows: 37.98 ± 29.27 kPa for control and 100.42 ± 79.84 kPa for T1DM mice cardiomyocytes (figure 6C).

**Figure 6.**
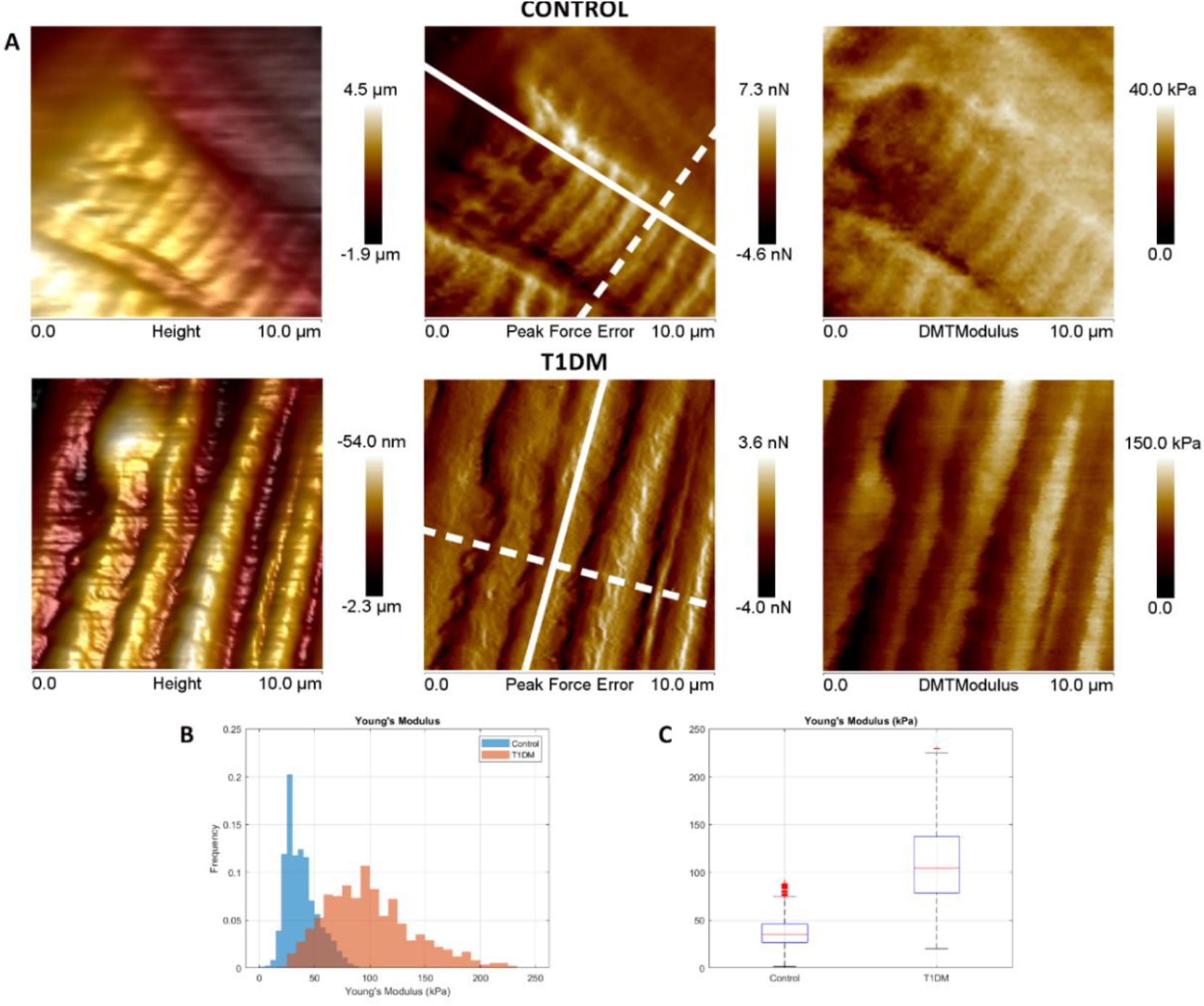
Topography and elasticity maps of living cardiomyocytes of control and T1DM mice, obtained by AFM, and statistical analysis. (A) Representative AFM topography images (3D height and peak force error) and elasticity map (DMT Modulus) of control (upper row) and T1DM (lower row) mice. Solid and dashed lines in peak force error images indicate long and short cardiomyocyte axes respectively. Control maps show two closely spaced parallel myofibrils that almost look like a single one. (B) Histogram showing retrieved values of Young’s Modulus from PF-QNM elasticity maps of cardiomyocytes of control and T1DM mice. (C) Elasticity maps of live T1DM cardiomyocytes were statistically different than control ones. Elasticity maps of T1DM were stiffer than the control. Mann-Whitney U and Kolmogorov-Smirnov tests (data expressed as medians ± SD) were used for comparing the data of control (n = 9) and T1DM (n = 8) mice (*p < 0.01).

## Discussion

Several findings highlight the complexity of the impact of diabetes on the myocardium^21,25-28^. Heart disease causes an alteration in myocardial relaxation and an increase in left ventricle stiffness, causing a decrease in ejection fraction^34,35^. The structural organisation of the cardiomyocyte is of utmost importance for its efficient functioning as a contractile unit in the heart. An essential component in mechanotransduction is the costamere complex, which is an ultraprotein structure made up of three complexes (dystroglycoprotein, integrin-talin-vinculin and spectrin complex). Their function is to connect the sarcolemma with the internal cytoskeleton. These complexes play a key role in the stability of the cell membrane and the transmission of force during contraction^1,36-38^. Alteration in any of these structural components can have a significant impact on cardiomyocyte function and ultimately overall cardiac function, underscoring the critical importance of understanding and maintaining structural organisation. It has been shown that force generation between cardiomyocytes cultivated and their substrate is mediated through the costamere^36^. Mechanical forces regulate the gain and loss of cell adhesion, stretching of the membrane and cytoskeleton, and cell compression due to changes in pressure^39^.

In this work, we found that some costamere proteins in the myocardium tissue and cardiomyocytes of T1DM mice had a significantly different spatial localisation than that of control mice. The results show that α5-integrin and vinculin proteins in control conditions have fewer clumps per μm^2^ than T1DM mice (Figures 2 and 4). Moreover, talin2 and vinculin proteins have more occupied areas in control samples than T1DM ones (Figures 3 and 4), This different spatial localisation of the costamere proteins could be affecting the mechanical properties of the myocardium. Another discovery of this work is that in most of the fibres of T1DM, it was very difficult to identify intercalated discs by transmitted light, and in those that could be identified, practically no signal from the presence of vinculin was observed. This result is reflected in the quantification of the number of intercalated discs per fibre (Figure 4). The observation is that vinculin in controls is located mainly in the intercalated discs, whereas in T1DM it has a more peripheral location along the myofibril. The cardiac intercalated disc is a specialised cell junction that mechanically and electrically couples cardiomyocytes. In many instances, a change in the structure of one protein is likely to significantly change its interaction with many other proteins^14,40-42^. Cardiac-specific knockout mice of vinculin lead to severely disrupted intercalated discs. Only 50% of mice survive past three months^6^. This decrease in vinculin in intercalated discs in T1DM cardiac tissue could indicate less communication and less cell adhesion, which could affect myocardial mechanics and cell communication. To rule out that the results were biased by the plane where the images were taken, Z-stacks were performed confirming that the results obtained do not have any bias.

The muscle is highly anisotropic, the transmission of biophysical force depends on the longitudinal or transverse direction of the force, as well as the balance between external and internal force generation^39^, and can occur parallel or lateral to the axis longitudinal of the sarcomere^43^. This could be due to the specific location and orientation of the mechanosensors, relative to the alignment of the cytoskeleton^44^. In the longitudinal direction, force is transduced from one sarcomere to the next within the same fibre until the end of the fibre is reached. Perpendicularly, lateral force transmission allows the transduction of a myofibril to a neighbouring myofibril until it reaches the costameric complex that channels the intracellular force through the sarcolemma to the extracellular matrix^43^. Likewise, just as the force generated by a cardiomyocyte can be transmitted longitudinally through cell-cell interactions, it can also be transmitted to adjacent cells laterally, through attachment to the extracellular matrix, allowing for coordinated contraction and relaxation of the functional syncytium and the myofibrillar apparatus of adjacent muscle cells^39^. It has been reported that longitudinal force transmission represents only 20-30% of the force generated by the sarcomeres, indicating that the main force vector is produced laterally. Hence, the location of the costamere makes it critical for its central role in the transmission of force^43^.

It has been reported that fibrotic and rigid environments destabilise cell-cell adhesions due to an increase in force generation, which stimulates the formation of focal adhesions, homologous to costameres^45^. Likewise, several cardiomyopathies, such as myocardial infarction, are characterised by increased fibrosis and cellular stiffness^45^, increased expression of integrins^46^, and lack of intercellular connections^47^, indicating that an increase in cell-extracellular matrix adhesions occurs while cell-cell adhesions decrease in cardiomyocytes^45^. It is important to highlight that the remodelling of intercellular adhesions could contribute to the redistribution of GAP junctions and the generation of arrhythmias, associated with many cardiomyopathies^47^. Our results are congruent with those that grew cardiomyocytes on a stiff matrix, which demonstrated an impaired contractile function with disarranged sarcomeres whereas cardiomyocytes grown on soft and medium matrices had highly organised sarcomeres^48^.

Our work also investigated whether these differences in the spatial arrangement of these proteins were due to differential protein expression in T1DM. Western blot analyses did not show these three proteins’ differential expression of T1DM compared with controls. These results are consistent with the work of Hersch et al., 2013^49^ where no costamere numbers were affected by substrate stiffness, and with the work of Romanelli et al.^21^ where a significant disordered arrangement of actin was observed in the diabetic myocardium without differences in the quantification of actin by Western blot. These results converge in the fact that diabetes produces changes at the level of the spatial organisation of several proteins in the cardiomyocyte although their expression remains constant.

Another essential component in this organisation is the T-tubule system, which consists of invaginations of the sarcolemma that extend into the interior of the cell. These tubules play a critical role in the transmission of electrical and mechanical signals, and calcium transport. The cardiac T tubular system provides structural integrity to the cardiac membrane during contraction/relaxation cycles^5,50^. Kostin et al. demonstrated an ordered organisation pattern in T-tubules. The cardiac T tubular system contains a subcellular scaffold that confers structural integrity to the cardiac T tubular membrane during contraction/relaxation cycles^5^. In the present work, it is evident how diabetes alters the orderly organisation of the T-tubules. Since α5-integrin is a transmembrane protein, the observation of α5-integrin in the middle plane of the cardiomyocytes, shows us that we are seeing α5-integrin located in the T-tubules. These results are according to Kostin et al.^5^ Comparing the distribution of α5-integrin between controls and diabetics, we can see that the controls maintain the orderly organisation of the T-tubules (Figure 2). In contrast, diabetic cardiomyocytes have a disorderly organisation. This fact is not minor since various works have already demonstrated the importance of T-tubules on calcium transport, and the latter with muscle contraction and relaxation^28,51^.

Cardiomyocytes are permanently exposed to mechanical stimulation due to cardiac contractility, hemodynamic pressure by blood, and elasticity/stiffness from the extracellular matrix. Changes in cardiac stiffness are hallmark characteristics of several cardiac diseases. Notwithstanding, we are just beginning to understand how biomechanical signalling changes in the diseased heart affect cardiomyocytes^13,52,53^. In 2014, Benech et al. observed a disordered alignment of myocardial cells, cell nuclei or irregular size and disordered myocardial fibre with an accumulation of interstitial collagen in samples of mice with diabetic hearts. In addition, the data obtained by using AFM demonstrated that diabetes had an impact on the nanomechanical properties of live cardiomyocytes, resulting in greater cellular stiffness in diabetic conditions. Furthermore, in that work, it was demonstrated that changes in cardiomyocyte stiffness were due, at least in part, to actin cytoskeleton dependence^28,31^. The maintenance of physiological cardiomyocyte stiffness determines proper cardiomyocyte functionality^53^. In 2021, Varela et al. obtained for the first time elasticity maps of cardiac cells in the H9c2 cell line^31^, but until now no elasticity maps of live diabetic primary cardiomyocytes had been obtained. Our work exposes, for the first time, elasticity maps of primary cardiomyocytes showing greater stiffness in T1DM cardiomyocytes than in control cardiomyocytes (according to Varela et al, 2021^31^). All the data obtained in the present investigation, added to the data obtained with the work of Romanelli et al.^21^, show intrinsic changes that T1DM causes in murine cardiomyocytes.

In summary, the present work shows, for the first time, experimental evidence that in the T1DM model used, intracellular changes occur related to cell-cell and cell-extracellular matrix communication which could be related to cardiac pathogenic mechanisms linked to the progress of diabetic cardiomyopathy. Our results demonstrate that T1DM induces changes in the spatial organisation of three costamere proteins (α5-integrin, talin-2 and vinculin), and in T-tubules organisation. These changes could contribute to alterations in cardiomyocytes’ mechanical and electrical properties and consequently of the myocardium, producing several cardiac pathologies. Finally, being able to understand the modifications that diabetes induces at the cellular level and being able to detect these alterations could in the future provide an early diagnosis of cardiac pathologies before they are evident at the organic level.

## Acknowledgements

We thank the IIBCE vivarium staff for the care and maintenance of the experimental animals.

